# Genetic interactions affect lung function in patients with systemic sclerosis

**DOI:** 10.1101/581553

**Authors:** Anna L. Tyler, J. Matthew Mahoney, Gregory W. Carter

## Abstract

Scleroderma, or systemic sclerosis (SSc), is an autoimmune disease characterized by progressive fibrosis of the skin and internal organs. The most common cause of death in people with SSc is lung disease, but the pathogenesis of lung disease in SSc is insufficiently understood to devise specific treatment strategies. Developing targeted treatments requires not only the identification of molecular processes involved in SSc-associated lung disease, but also understanding of how these processes interact to drive pathology. One potentially powerful approach is to identify alleles that interact genetically to influence lung outcomes in patients with SSc. Analysis of interactions, rather than individual allele effects, has the potential to delineate molecular interactions that are important in SSc-related lung pathology. However, detecting genetic interactions, or epistasis, in human cohorts is challenging. Large numbers of variants with low minor allele frequencies, paired with heterogeneous disease presentation, reduce power to detect epistasis. Here we present an analysis that increases power to detect epistasis in human genome-wide association studies (GWAS). We tested for genetic interactions influencing lung function and autoantibody status in a cohort of 416 SSc patients. Using Matrix Epistasis to filter SNPs followed by the Combined Analysis of Pleiotropy and Epistasis (CAPE), we identified a network of interacting alleles influencing lung function in patients with SSc. In particular, we identified a three-gene network comprising *WNT5A, RBMS3*, and *MSI2*, which in combination influenced multiple pulmonary pathology measures. The associations of these genes with lung outcomes in SSc are novel and high-confidence. Furthermore, gene coexpression analysis suggested that the interactions we identified are tissue-specific, thus differentiating SSc-related pathogenic processes in lung from those in skin.

**Author summary:** Systemic sclerosis (SSc), or scleroderma, is a devastating autoimmune disease. Patients experience progressive fibrosis of their skin and internal organs, reduced quality of life, and increased risk of death. Lung disease associated with SSc is particularly dangerous and is currently the leading cause of death in SSc patients. There are no specific treatments for SSc or SSc-related lung disease, but promising work in the genetics of this disease has identified more than 200 genetic variants that influence SSc [1]. Piecing together how genetic variants interact with each other to influence disease may provide clues for targeted therapies. Here we present a novel analytical approach for identifying genetic interactions in a human disease cohort. In this approach we first filtered SNPs to those that are most likely to interact to influence the disease traits. We then applied the Combined Analysis of Pleiotropy and Epistasis (CAPE), which combines information across multiple traits to increase power to detect genetic interactions. Using this approach, we identified a three-gene network among *MSI2, WNT5A*, and *RBMS3* that influenced autoantibody status and lung function in a cohort of 416 SSc patients. Gene expression data suggest that this interaction network is tissue- and disease-specific, and may thus provide a specific target for SSc therapy.

## Introduction

Systemic sclerosis (SSc), or scleroderma, is a complex autoimmune disease associated with significant morbidity and mortality. It is characterized by widespread fibrosis of skin and internal organs, as well as vasculopathy [2]. The most common cause of death in SSc is lung disease [3]. Roughly a quarter of people with SSc develop interstitial lung disease, which involves lung fibrosis, vascular hyperreactivity, and inflammation [4]. Another 7-13% of patients develop pulmonary arterial hypertension, which is characterized by vascular injury and occlusion, vasoconstriction, and dysregulated angiogenesis [4]. Both conditions lead to reduced lung function and increased risk of death. The pathogenesis of lung disease in SSc is not understood well enough for development of specific treatments, and current treatments rely primarily on non-specific immune suppression [5]. There is a need to identify new molecular drivers of lung disease in SSc, as well as how these drivers interact with other genes to influence pathogenesis.

A standard approach to discovering molecular drivers of lung disease in SSc is to identify genetic variants associated with lung outcomes. Genetic studies have been tremendously successful in identifying genetic variants associated with SSc and its complications. In a reflection of the complexity of the disease, variants in over 200 genes have been implicated in SSc risk and progression [1], which has greatly increased our understanding of the development of SSc [6-8] and may aid in personalized disease monitoring and treatment [9]. The next step in this line of inquiry is to incorporate genetic complexity into models that determine how variants interact with each other to influence disease. By explicitly modeling genetic interactions, or epistasis, we can build understanding of how molecular pathways work in concert to drive SSc pathology.

Initial studies of genetic interactions in SSc have been promising. Epistasis between polymorphisms in the HLA region and cytokines has been shown to predict SSc risk [10], development of severe ventilatory restriction [11], and digital ulcer formation [12] in SSc patients. However, progress in this search is limited by a number of challenges. The rarity of the disease and its clinical heterogeneity add to difficulties present in all human genetic studies, such as low minor allele frequencies and the large number of potentially relevant variants. Non-parametric tests such as Multifactor Dimensionality Reduction (MDR) [13] have been successful in identifying the interactions that have been identified thus far [10-12]. These findings suggest additional, complementary interaction analyses may further dissect the genetic complexity of SSc and other common diseases.

Here we present a novel approach that increases the power to detect genetic interactions in human genome-wide association studies (GWAS). We previously developed the Combined Analysis of Pleiotropy and Epistasis (CAPE) to model epistatic interactions [14, 15]. CAPE increases power to detect and interpret genetic interactions by combining information across multiple traits into one consistent model. We have demonstrated its ability to identify novel genetic interactions not detectable by other methods [24, 26]. For this study, we combined CAPE with a filtering step, which filtered the SNPs to those most likely to be involved in genetic interactions. We used Matrix Epistasis [16], an ultra-fast method for exhaustively testing epistasis in genome-wide SNP data. Candidate SNP pairs were then analyzed with CAPE and significance was assessed with permutation tests.

We applied this approach to genetic and clinical data from a cohort of patients with SSc (dbGaP accession phs000357.v2.p1). To capture aspects of lung disease and autoimmunity, we analyzed two measures of lung function, forced vital capacity (FVC) and diffusion lung capacity (DLC), as well as two autoantibody staining patterns: centromeric, and nucleolar. Anti-centromere autoantibodies (ACA) are associated with pulmonary hypertension, and anti-nucleolar autoantibodies (ANA) are associated with progressive interstitial lung disease and pulmonary arterial hypertension [17]. Thus, there may be common genetic pathways underlying both autoantibody status and lung function that we can identify by analyzing all four traits simultaneously.

## Results

### Trait Descriptions

We analyzed a cohort of 416 patients with lung function and autoantibody measurements (369 females and 47 males). The two lung traits, forced vital capacity (FVCP) and diffusion lung capacity (DLCP), were measured as a percentage of the total predicted by patient demographic parameters, such as age and weight [18]. Both traits were roughly normally distributed, with maximum values greater than 100%, and the traits were modestly correlated (Pearson’s *r* = 0.48, *p* = 1.3×10^-25^ (Fig 1A). We used rank Z normalization on both traits before performing association tests. The distribution of autoantibodies is shown in Fig 1A. About half of patients were negative for both autoantibodies, while the other half were positive for one or the other. Only five patients were positive for both autoantibodies.

**Fig 1.**
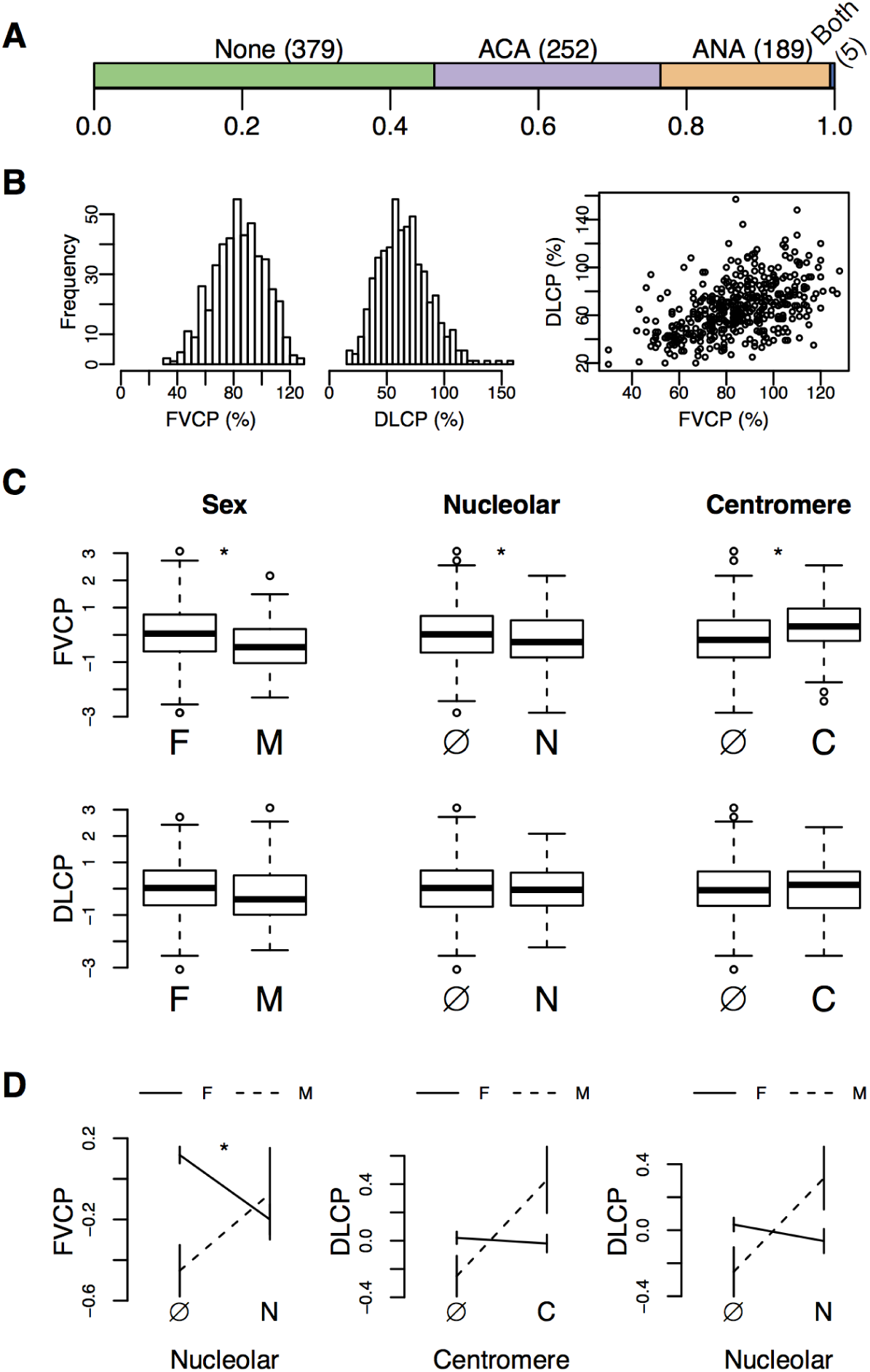
Trait distributions and correlations. (A) Colored bars show the proportion of patients that were negative for both autoantibodies (None), positive for either centromere (ACA) or nucleolar (ANA) autoantibodies, or positive for both (Both). (B) Histograms showing the distribution of FVCP (%) and DLCP (%). The distributions are roughly normal. The third panel shows the correlation between FVCP and DLCP. They were moderately correlated (*r* = 0.48, *p* = 1.3×10^-25^). (C) Box plots show associations between lung traits and binary traits. Sex, nucleolar status, and centromere status all had significant effects on FVCP. None of the binary traits were significantly associated with DLCP. Significant effects (Student’s t test *p* < 0.05) are identified with an asterisk (*). Autoantibody status is denoted as ANA (N), ACA (C) or absent (Ø). (D) These plots show interactions between binary traits that affect the lung traits. Sex and nucleolar status interacted significantly to influence FVCP. Interactions between sex and autoantibody status on DLCP were not quite significant, but trended toward significance (*p ≈* 0.06).

FVCP was influenced by sex and autoantibody status (Fig 1B). Males had significantly lower FVCP on average than females (Student’s t test *p* = 0.0063). The presence of the two autoantibodies had opposite effects. The presence of ANA significantly reduced FVCP (Student’s t test *p* = 0.033), while the presence of ACA significantly improved FVCP (Student’s t test *p* = 5.1*×*10^-7^). Neither autoantibody nor sex had any significant association with DLCP (all Student’s t test *p* > 0.05).

We also looked for interactions between the autoantibodies and sex that influenced the lung traits (Fig 1C). Presence of ANA and sex interacted nominally significantly to influence FVCP (*p* = 0.042). In this interaction, females positive for ANA had lower FVCP on average than females negative for ANA. However, in males, the effect was the opposite. Males positive for ANA had higher FVCP on average than males negative for ANA. None of the other factor combinations had any significant effects on either FVCP or DLCP (all Student’s t test *p* > 0.05); however sex and autoantibody status trended toward a significant interaction for DLCP. The presence of each autoantibody had very little effect on DLCP in females, but increased DLCP in males.

### Single-locus association identified four SNPs with significant main effects on ACA

The patients in this study were genotyped at 601,273 SNPs. However, because detection of genetic interactions can be confounded by low minor allele frequencies (MAF), we first filtered this data set to 243,662 SNPs with MAF≥0.1. We then performed all single marker regressions (Methods). The purpose of this analysis was two-fold. First it allowed us to determine whether the number of patients in this cohort was sufficient to recapitulate known associations between SNPs and SSc traits. Second, we used the *p* value distributions from this analysis to investigate whether population structure in this cohort might affect the SNP associations (Methods). There was no systematic effect of population structure in this cohort (Fig S1). We therefore did not use a correction for population structure in this analysis.

The SNP association tests identified significant main effects in the HLA region influencing the presence of ACA (Table 1). One of these SNPs (rs9275390) is associated with HLA-DBQ1 and was previously associated with positive ACA status in an SSc GWAS [36]. Another SNP, rs660895, is associated with HLA-DRB1, and was previously associated with risk of the autoimmune disease Rheumatoid Arthritis [37].

**Table 1.** Information about position of SNPs significantly associated with ACA (*p* < 0.01).

| SNP | Chr | Position | Nearest Gene | Annotation | Distance |
| --- | --- | --- | --- | --- | --- |
| rs4248166 | chr6 | 32398644 | BTNL2 | intronic |  |
| rs660895 | chr6 | 32609603 | HLA-DRB1 | upstream | 19755 |
| rs7755224 | chr6 | 32684540 | HLA-DQB1 | upstream | 16157 |
| rs9275390 | chr6 | 32701379 | HLA-DQB1 | upstream | 32996 |

### CAPE identified multiple epistatic interactions influencing lung and autoantibody traits

We next performed pairwise SNP associations to identify interactions between SNPs. As CAPE is relatively computationally intensive, performing exhaustive pairwise testing (> 1.8×10^11^ tests) was not feasible. We thus filtered the SNPs to 1500 that were likely to participate in epistatic interactions influencing more than one trait (Methods). In addition, we decomposed the traits into orthogonal eigentraits (ETs) using singular value decomposition (SVD) (Fig 2A) (Methods). Traits that are moderately correlated may share common underlying biological processes. Using ETs concentrates signals from these underlying processes into single traits thereby increasing power to map them genetically. We selected the first two ETs for analysis. Each ET describes the contrast between one autoantibody and the two lung traits. After calculating CAPE coefficients associated with the two ETs, we rotate the coefficients back into trait space. CAPE thus identifies interactions associated with all traits simultaneously. By selecting only two ETs we lose some biological information contained in the other two ETs, but we simultaneously gain power to identify interactions influencing the biological processes encoded by the first two eigentraits.

**Fig 2.**
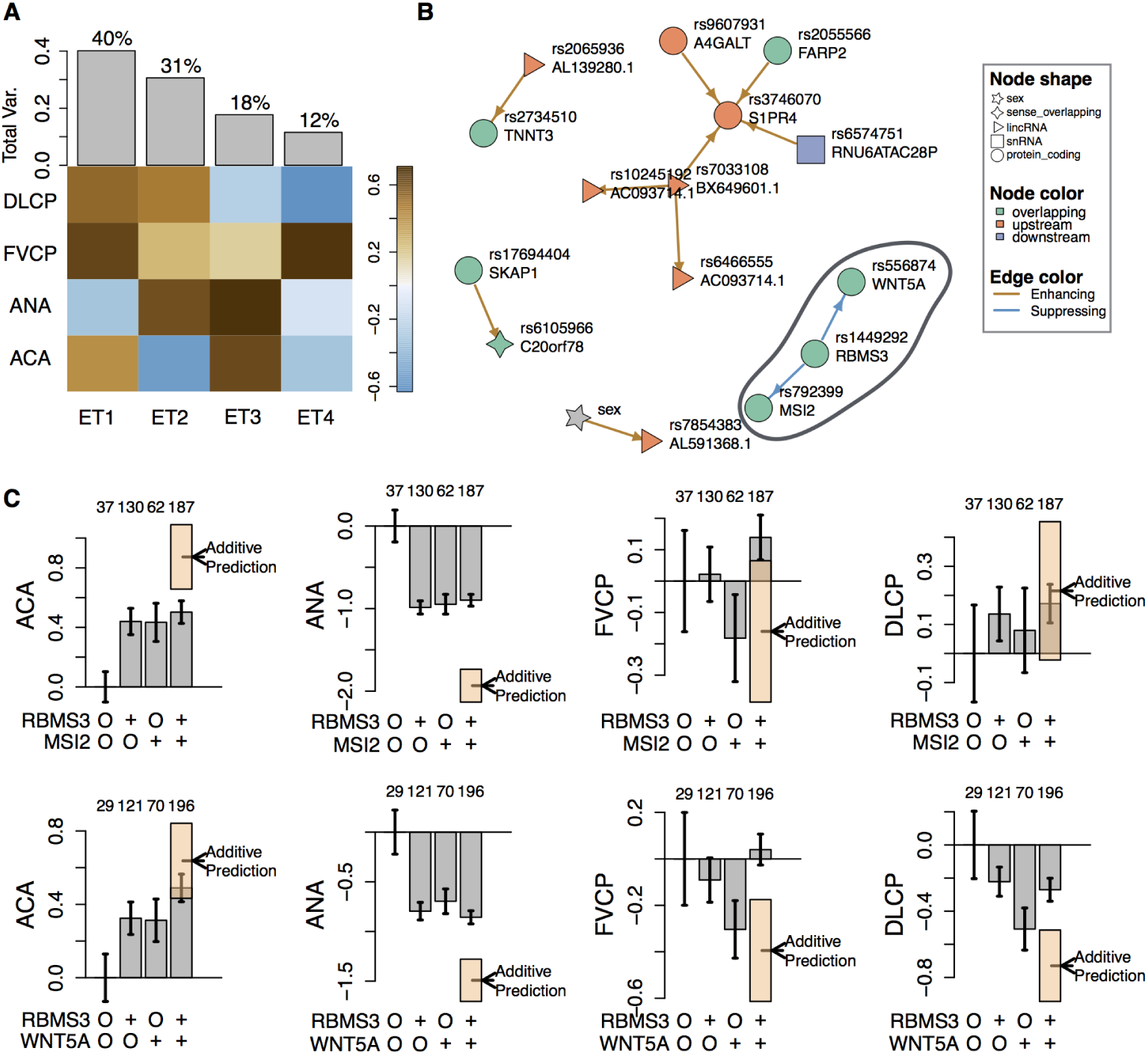
Genetic interactions influence lung function and autoantibodies in SSc. (A) The relative contributions of each trait to each eigentrait (ET). Gray bars indicate how much overall variance each ET captures. We used the first two ETs, each of which contrasts one of the autoantibodies, with the other autoantibody and the lung function traits. These two ETs capture 71% of the overall trait variance. (B) The network of significant SNP-SNP interactions discovered by CAPE. Each node in the network represents a SNP. Each is labeled with the rs number, as well as with the name of the nearest gene. The color of each node indicates whether the SNP is upstream, downstream, or overlapping the gene. The shape of each node indicates the type of gene, protein coding, lncRNA, etc. The links between the nodes are colored to indicate whether each significant interaction is enhancing or suppressing. Three SNPs (rs556874, rs1449292, and rs792399) form a three-node subnetwork we called the Wnt subnetwork (circled). Each of these SNPs overlaps a protein coding gene. (C) The effects of the Wnt subnetwork interactions on the four traits. The x axis in each plot indicates the four possible genotypes for each set of two SNPs. SNPs are labeled by their overlapping gene for clarity. The “O” indicates the reference genotype, and “+” indicates the alternate genotype. Gray bars indicate the mean trait value for the group of patients with the genotype indicated on the x-axis. The number above each bar indicates how many patients are in that group. Error bars show standard error. The additive prediction for each double alternate genotype is indicated by the dashed line and the error of the estimate is shown in the orange box. Gray bars that fall outside the orange box indicate significant interactions between the alleles. In A and C traits are abbreviated as follows: percent predicted forced vital capacity (FVCP) and percent predicted diffusion lung capacity (DLCP), antinucleolar autoantibodies (ANA), anticentromere autoantibodies (ACA).

We found a network comprising 11 directed genetic interactions involving 15 SNPs and the covariate sex (Fig 2B). The two most significant interactions (largest standardized effects) formed a single subnetwork among SNPs rs556874, rs1449292, and rs792399 (Fig 2B). This subnetwork was notable in that, not only did it contain the most significant interactions, but all three SNPs were intronic in protein coding genes *WNT5A, RBMS3*, and *MSI2*. The SNP rs556874 is located in the fourth intron of *WNT5A*, which is a member of the large family of Wnt ligands. Wnt signaling is a widely important family of signaling pathways integral to embryonic development, carcinogenesis, and many other processes. The SNP rs1449292 is located in an intron of *RBMS3*. This gene encodes a member of a small family of proteins that bind single-stranded RNA and DNA to regulate a wide range of biological processes. The SNP rs792399 is located in an intron of *MSI2* about which little is known. We focused on this subnetwork, which we refer to as the “Wnt subnetwork,” for the remainder of the analysis.

### Interactions in the Wnt subnetwork were less than additive

The interactions in the Wnt subnetwork were suppressing (Fig 2A). This means that in the presence of the alternate allele at one locus, the phenotypic effects of the interacting locus were suppressed. In this case, the suppressing interactions resulted less-than-additive interactions. These effects can be seen in Fig 2C. The *RBMS3*-*MSI2* interaction primarily affected the two autoantibody traits. The alternate alleles of the SNPs each had a negative effect on autoantibody presence. However, the effects of both alternate alleles together was redundant and did not decrease autoantibody presence beyond either of the individual alleles. The effects of this interaction on lung function were not different from additive.

The *RBMS3*-*WNT5A* interaction affected both lung function traits and ANA. The effects on ANA were similar to the effects of the *RBMS3*-*MSI2* interaction: individual alternate alleles decreased ANA incidence, but having two alternate alleles did not further decrease incidence beyond that seen for a single alternate allele. For both lung function traits, the *WNT5A* alternate allele had a large negative effect in the presence of the *RBMS3* reference allele, but this effect was completely suppressed in the presence of the *RBMS3* alternate allele.

### Expression of *RBMS3* and *WNT5A* was dysregulated in SSc lung

We investigated whether the candidate genes in the Wnt subnetwork were dysregulated in SSc patient lung tissue (Methods). *WNT5A* (Student’s t test *p* = 0.02) and *RBMS3* (Student’s t test *p* = 0.011) were both significantly upregulated in SSc lung tissue relative to healthy lung tissue (Fig 3A), supporting the hypothesis that the activity of genes we identified genetically are aberrantly regulated in SSc. SNPs affecting the expression of these genes may also affect SSc presentation. We could not assess the differential expression of *MSI2*, as it was not present in the data set.

**Fig 3.**
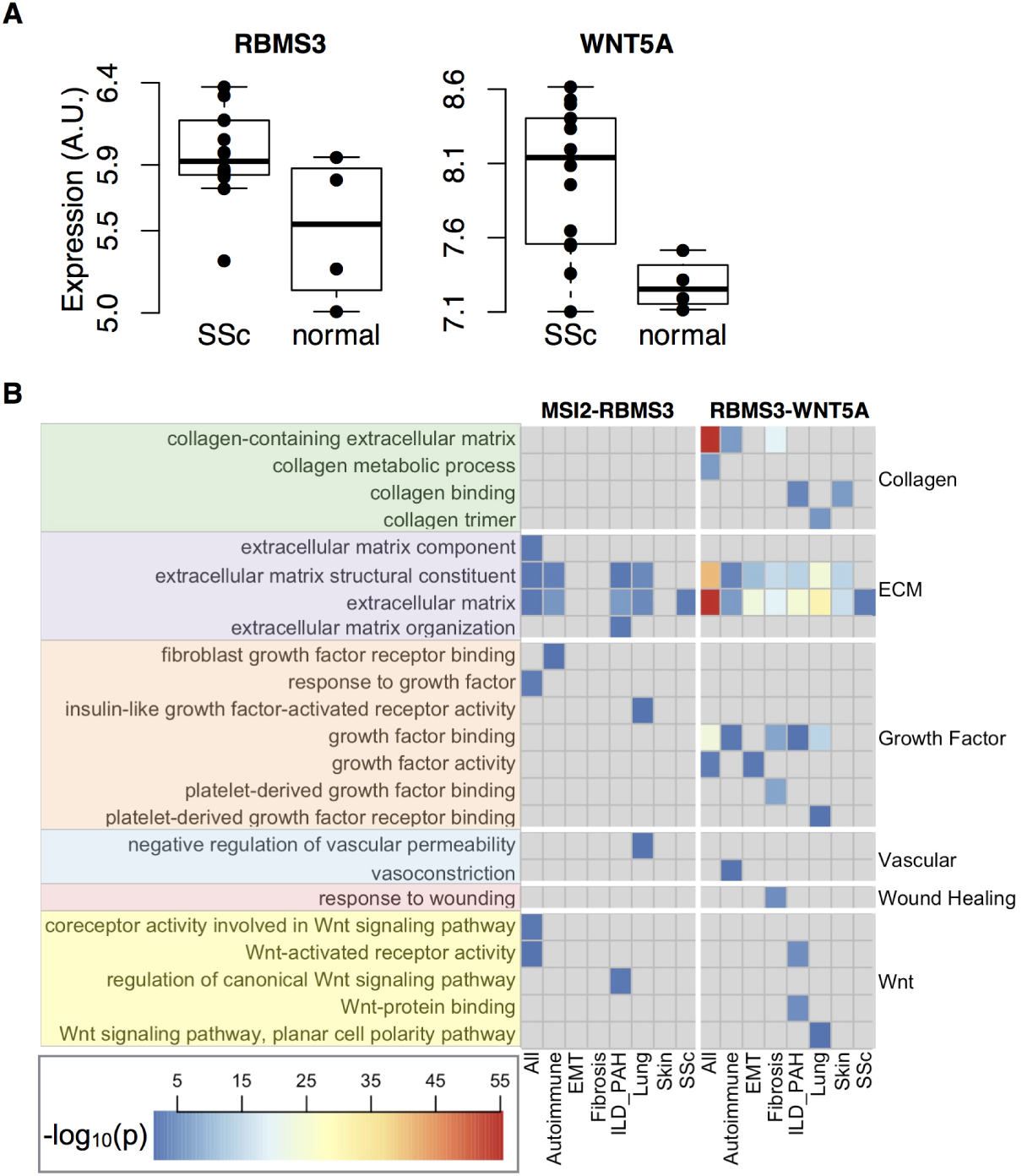
Expression of subnetwork genes in SSc. (A) Both *WNT5A* (Student’s t test *p* = 0.011) and *RBMS3* (Student’s t test *p* = 0.02) were significantly overexpressed in lung tissue from SSc patients with interstitial lung disease. (B) The –*log*_10_ (*p*) for enriched GO terms in gene sets derived from SEEK (Methods). Each column shows results from a subset of the data sets on GEO. They are as follows: All - all data sets on GEO, Autoimmune - data sets studying autoimmunity, EMT - data sets studying epithelial to mesenchymal transition, Fibrosis - data sets studying fibrosis, ILD PAH - data sets studying interstitial lung disease (ILD) or pulmonary hypertension (PAH), Lung - data sets using lung tissue, Skin - data sets using skin tissue, SSc - data sets studying SSc. Each row shows the results for one GO term identified on the right-hand side. The GO terms are grouped functionally into six groups: collagen (green), extracellular matrix (purple), growth factor (orange), vascular (blue), wounding, (red), and Wnt signaling (yellow). The gene names at the top of the figure indicate that the left hand side of the figure shows results for the *RBMS3*-*MSI2* interaction, and the right side shows results for the *RBMS3*-*WNT5A* interaction. The legend indicates how the colors of the cells relate to the *–log*_10_(*p*) of the term enrichment.

### Co-expressed genes were functionally enriched for processes related to SSc

To further investigate biological processes affected by the Wnt subnetwork, we searched across GEO for genes that were co-regulated with the gene pairs in the Wnt subnetwork. Genes with correlated expression often function together, and co-expression is frequently used to elucidate gene function [34, 38-40]. We used this principle to identify possible functional roles of the interactions in the Wnt subnetwork. To investigate whether the interactions might be context-specific, we searched across multiple subsets of data sets in GEO (Methods). For example, to identify lung-specific functions of each gene pair, we searched for genes with correlated expression among data sets that were obtained from lung tissue. To identify functions that were related to autoimmunity, we searched only among data sets that analyzed autoimmune disease. All data set subsets are listed in (Table 2).

**Table 2.**
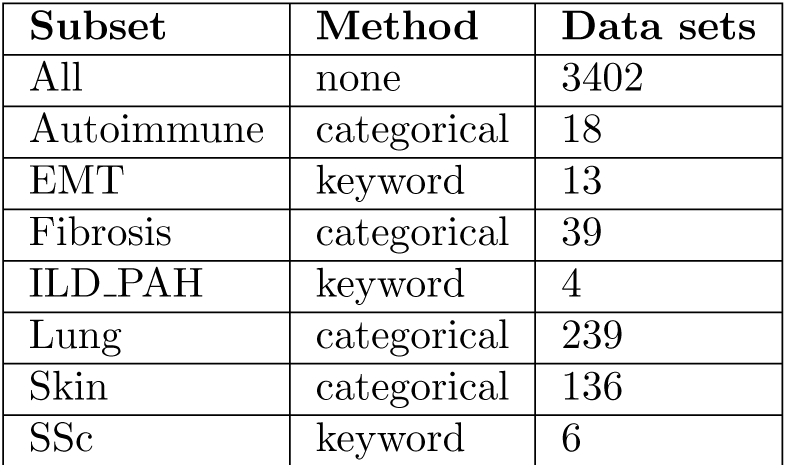
The terms used to refine searches across gene expression data sets in GEO. Subsetting was performed either by selecting a pre-defined subset (categorical) or by using key words (keyword) (Methods). The final column indicates how many data sets were included when subset criteria were applied.

| Subset | Method | Data sets |
| --- | --- | --- |
| All | none | 3402 |
| Autoimmune | categorical | 18 |
| EMT | keyword | 13 |
| Fibrosis | categorical | 39 |
| ILD_PAH | keyword | 4 |
| Lung | categorical | 239 |
| Skin | categorical | 136 |
| SSc | keyword | 6 |

In each of these data set subsets, we queried the *WNT5A*-*RBMS3* gene pair separately from the *RBMS3*-*MSI2* gene pair. For each query, we identified sets of genes whose expression was significantly correlated with the two query genes (*p* < 0.05), and used the R package gProfileR [35] to look for functionally enriched GO terms in each gene set. Multiple GO terms were significantly associated with the gene set from each query. For a complete list of the enriched terms, see Supplementary Fig 1. To focus on terms that are strongly related to SSc, we only analyzed GO terms that included the following SSc-related strings: “collagen”, “matrix”, “fibroblast”, “growth factor”, “vascular”, “vaso”, “wound”, and “wnt” (Methods). These terms, and their enrichment *p* values across all queries are shown in Fig 3B.

Both gene pairs were significantly correlated with extracellular matrix (ECM) GO terms across multiple tissue types and conditions (Fig 3); however, only the *RBMS3*-*WNT5A* pair was significantly correlated with genes involved in collagen indicating that this pair may affect ECM through effects on collagen, whereas the *RBMS3*-*MSI2* pair may influence ECM in a collagen-independent manner. The *RBMS3*-*WNT5A* pair also had multiple enrichments relating to growth factor activity, particularly in lung tissue, and in lung disease (ILD PAH). This suggests that the pair may be involved in growth factor signaling in a tissue-specific manner. All significant enrichments for the *RBMS3*-*MSI2* interaction were in lung, lung disease, and autoimmune conditions, suggesting that this interaction may be specific to lung tissue, and may be aberrantly regulated in autoimmune conditions. The *RBMS3*-*WNT5A* interaction had significant enrichments in the data sets relating to fibrosis and EMT, while the *RBMS3*-*MSI2* interaction did not, suggesting that the *RBMS3*-*WNT5A* interaction may be more functionally related to these processes than the *RBMS3*-*MSI2* interaction.

## Discussion

We used a novel analytical approach to identify an epistatic interaction network influencing lung function in patients with SSc. In this approach we used three strategies to increase power to detect interactions in a patient cohort. First, we used a SNP filtering step, to select the most highly epistatic and pleiotropic SNPs for further testing. This step not only identified SNPs that were most likely to interact, but also substantially reduced the number of statistical tests. Second, we combined information across multiple related traits: two autoantibody traits, and two lung function traits. And third, we decomposed the traits into orthogonal eigentraits and used only the top two eigentraits in our study. This step reduced noise and captured mappable signals that related to all four traits. Thus, using Matrix Epistasis to pre-filter SNPs combined with CAPE allowed us to identify high-confidence genetic interactions in a relatively small human cohort.

The resulting genetic interaction network contained a notable subnetwork consisting of two interactions between three SNPs. Because the interactions in this network were highly significant and because all three SNPs were located within gene bodies, we focused on this subnetwork for further analysis. The genes tagged by this subnetwork were *WNT5A, RBMS3*, and *MSI2*, and we refer to the subnetwork as the Wnt subnetwork. None of the three genes in the Wnt subnetwork had previously been associated with lung function in SSc, although they have all been implicated in lung disease, as well as in molecular processes that are known to be involved in SSc pathogenesis.

*WNT5A*, and Wnt signaling more generally, have been implicated in many processes known to be involved in SSc pathogenesis [41-44], such as angiogenesis [45], keratinocyte differentiation and inflammation signaling [46], and fibroblast proliferation [47]. *WNT5A* has also been linked to cellular transdifferentiation, a family of processes by which differentiated cell types dedifferentiate and redifferentiate into another cell type. Transdifferentiation processes are critical for development, wound healing and tissue regeneration [48]. However, when they are dysregulated, these processes contribute to metastases in cancer, and widespread fibrosis seen in fibrotic diseases [48]. There are three types of transdifferentiation that have been linked to fibrosis in SSc: epithelial to mesenchymal transition (EMT) [49], endothelial to mesenchymal transition (EndoMT) [50] and adipocyte-myofibroblast transition (AMT) [51].

Each of these processes has been hypothesized to be a source of excess myofibroblasts in SSc tissues. Myofibroblasts differentiate from fibroblasts in response to signaling from TGF-*β*, as well as other growth factors and cytokines. In healthy individuals, myofibroblasts play an important role in wound healing. They contract to close wounds and deposit ECM to seal wounds and to provide a scaffold for the re-establishment of epithelial tissue. In SSc and other fibrotic disease however, there are increased numbers of myofibroblasts with a concomitant increase in the deposition of extracellular matrix, which causes tissue stiffening [49, 52]. EMT has been directly implicated in lung fibrosis in mice [53, 54], and may be an important source of myofibroblasts in SSc skin [52]. Wnt signaling, and *WNT5A* in particular, has been shown to mediate EMT in melanoma [55] and lung cancer [56].

Endothelial to mesenchymal transition (EndoMT) has also been proposed to be a source of myofibroblasts in SSc [50, 57, 58]. Originally thought only to occur in developing embryos, EndoMT has been demonstrated to occur in adult cattle [59] and mice [60]. TGF-*β*-induced EndoMT has since been shown to be involved in the bleomycin model of pulmonary fibrosis as well as SSc-associated pulmonary hypertension [58, 61]. Furthermore, there is evidence to suggest that Wnt signaling may mediate EndoMT [58].

The third transdifferentiation process to be proposed as a source of myofibroblasts in SSc is AMT [51]. Marangoni et al. (2015) showed that when adipose-derived stem cells from healthy donors were incubated with TGF-*β*, they lost markers of adipocytes, and gained markers of myofibroblasts. In this experiment *WNT5A* was strongly upregulated in the transitioning cells.

Like *WNT5A, RBMS3* has been linked to transdifferentiation. Upregulation of *RBMS3* inhibits EMT in lung cancer through the inhibition of canonical Wnt signaling [62, 63]. By downregulating MMP2 and *β*-catenin, *RBMS3* also inhibits microvessel formation [64]. Loss of microvessels and reduced angiogenesis are hallmarks of SSc [65, 66]. *RBMS3* may also directly influence collagen synthesis [67, 68], which is highly upregulated in SSc. Although *RBMS3* has not been directly linked to SSc, SNPs in *RBMS3* have been associated with risk of another autoimmune disease affecting connective tissue called Sjögren’s syndrome [69].

Of the three top candidate genes, the least is known about *MSI2*, and it is most studied in the context of cancer and its influence on EMT [70, 71]. *MSI2* is upregulated in metastatic-competent lung cancer cell lines, and aggressively metastatic patient tumors have upregulated *MSI2* [71]. Conversely, depletion of *MSI2* resulted in reduced metastatic potential [71]. As mentioned previously, EMT may be an important process in SSc pathogenesis. *MSI2* is also related to Wnt signaling and may influence hepatocellular carcinoma outcomes through dysregulation of Wnt signaling [72].

Thus, all three genes in the Wnt subnetwork have been implicated in mediating the transdifferentiation process of epithelial to mesenchymal transition (EMT). Both EMT [73] and EndoMT [58] are known to take place in the lung and have demonstrated roles in SSc [49, 61]. These results suggest that EMT, or another similar transdifferentiation, may play a role in lung disease pathogenesis and lung function in SSc patients, and that genes in the Wnt subnetwork we identified here may be involved.

Furthermore, all three genes in the Wnt subnetwork have been shown to be associated with Wnt signaling. These pathways are known to be important both in lung development, and in lung fibrosis in SSc [74, 75]. Wnt signaling, both canonical (*β*-catenin mediated) and non-canonical (not *β*-catenin mediated), has been shown to drive EMT. Upregulation of *WNT5A* increases EMT in lung cancer [56] possibly through stimulation of the non-canonical PKC Wnt pathway [55]. Similarly, upregulation of *MSI2* is an indicator of increased metastatic potential in lung cancer [71] and hepatocellular carcinoma [71], and is proposed to stimulate EMT through stimulation of canonical Wnt signaling [72]. *RBMS3* has been shown to reduce EMT and metastatic capacity of lung cancer cells [63] and reduces EMT in gastric cancer cells through suppression of canonical Wnt signaling [62]. Taken together, these results suggest that the Wnt subnetwork we identified here may mediate cellular transdifferentiation in the SSc lung through regulation of Wnt signaling.

Importantly, the Wnt subnetwork includes not only three novel candidate genes, but also the genetic interactions between them. These interactions suggest functional relatedness, which is broadly supported through connections in the literature with EMT and Wnt signaling. But we can also make more direct connections between the specific genes by using a more agnostic gene expression-based approach. We looked at genes that were co-expressed with each pair of subnetwork genes across multiple different tissue and disease contexts, including lung tissue, lung disease, and skin tissue. We found that the gene pairs were associated with multiple processes related to SSc pathogenesis, such as ECM organization, growth factor response, vascular abnormalities, and Wnt signaling. By comparing the enrichments related to the two interactions, we can generate hypotheses about their specific involvement. For example, both gene pairs were co-regulated with genes involved in ECM-related GO processes; however only genes co-expresed with *RBMS3* and *WNT5A* were associated with collagen, suggesting that *RBMS3* and *MSI2* together may influence a distinct aspect of ECM. Interestingly, functional enrichment of co-expressed genes also appeared to be context-specific. SSc-related functional enrichments were seen predominantly in the lung and lung disease (ILD PAH) data sets, as well as in autoimmunity data sets, but not in skin. We interpret this to mean that both genetic interactions may be tissue- and disease-specific, and that they were only detectable because we looked specifically at lung function outcomes in an SSc cohort. Thus, the interactions we identified may not be related to skin pathology. Validation experiments, therefore, should focus on lung tissue.

We further hypothesize that these interactions may be particularly important to lung function in a disease state. All three SNPs we identified have high minor allele frequencies in the general population (rs556874 MAF≈0.4, rs1449292 MAF≈0.5, and rs792399 MAF≈0.4) [76]. We hypothesize that these SNPs are unlikely to have any measurable effect on lung function in healthy individuals, but that the interactions between them become important in disease. We further hypothesize that because these alleles are so common, they may not increase the risk of developing SSc, but that they affect the course of the disease once established. Importantly, we performed our genetic analysis on a cohort of patients with SSc that did not include controls. Although a cohort containing only affected individuals may have few individuals, thereby reducing power to detect interactions, we assert that this type of case-only study is important in detecting disease-specific interactions that would not be detectable in a data set that included controls.

In this study we have taken a systems approach to identifying potential SSc-related processes by analyzing genetic interactions that influence multiple disease traits simultaneously. By addressing the complexity of this disease specifically in our models we increased power to detect novel interactions among highly promising candidate genes. The interactions, furthermore, may be disease- and tissue-specific. Identifying such specific interactions may provide a promising avenue forward in drug target discovery. The genes identified in this study are all widely expressed, and targeting them broadly may result in unintended effects. However, because evidence suggests that the interactions between the genes are tissue- and disease-specific, perhaps the interactions themselves are better targets than the genes. For example, rather than attempting to reduce *WNT5A* expression globally, a specific targeting of its interaction with *RBMS3* may provide a novel approach to treating SSc-related lung disease without widespread off-target effects. Known genetic interactions may also help predict disease course in individual SSc patients. If it is known, for example, that a patient has the reference *RBMS3* allele and the alternate *WNT5A* allele, this might indicate a highly likelihood of developing severe SSc-related lung disease, and testing and treatment of lung disease could be started early and aggressively. Genetic interactions offer a rich source of information for the understanding and treatment of disease, and new computational methods that take genetic and physiological complexity into account have the potential to build on the past success of GWAS and to piece together connections between individual findings that are currently isolated from one another.

## 1 Materials and Methods

### Methods workflow

All code used for computational analyses in this paper are available in a supplemental workflow available at https://github.com/annaLtyler/SSc_Epistasis_Workflow.

### Genetic and clinical data

Genetic data and autoantibody status were obtained from the database of Genotypes and Phenotypes (dbGaP) Accession phs000357.v1.p1.

The full data comprised 833 SSc patients, including 741 females and 85 males. Genotypes were measured at 601,273 SNPs using the Human610 Quadv1 B platform. Patients were assessed for the presence of multiple autoantibodies and lung function tests were performed. Values for autoantibody phenotypes were 1 (present) and 0 (absent), and values for test of lung function included forced vital capacity and diffusion lung capacity. Both measurements were represented as the percent of the predicted value based on factors such as age and sex [18].

We analyzed two lung function traits, percent predicted forced vital capacity (FVCP) and percent predicted diffusion lung capacity (DLCP) as well as two autoantibody patterns, anti-centromere (ACA) and anti-nucleolar (ANA). There were 416 patients (369 females and 47 males) with measurements for all four traits, and all subsequent analyses include only these patients.

### Expression data

We obtained gene expression data from the Gene Expression Omnibus (GEO) [19, 20]. The data set we used (accession number GSE76808) compared gene expression in lung biopsies taken from patients with SSc and interstitial lung disease (ILD) to biopsies of healthy tissue from lung cancer patients undergoing surgical resection [21]. The available data were *log*_2_ normalized, and we did not perform further normalization. Group-wise comparisons were performed with Student’s two-sample *t*-tests.

### SNP Filtering

We first reduced to 243,662 SNPs with minor allele frequency (MAF) ≥0.1, a relatively high cutoff due to our pair-wise testing strategy. Because analysis of all pairwise combinations of the filtered SNPs was computationally infeasible, we further filtered the SNPs to the top 1500 SNPs that were most likely to participate in genetic interactions influencing the four traits. For this filter we used Matrix Epistasis [16], an ultra-fast method of calculating interactions between SNPs. Using Matrix Epistasis, we were able to calculate interaction scores for all SNP pairs exhaustively for each of the four traits. We then used an iterative selection scheme to identify SNPs that were most epistatic and most pleiotropic (affecting more than one trait). In the first stage of the search, we identified SNP pairs that interacted significantly to influence more than one trait. There were 52 pairs of SNPs interacted significantly (*p* < 1×10^-6^) to affect both lung traits. No other SNP pairs influences more than one trait. In the next stage of the search, we looked for individual SNPs that were in epistatic interactions affecting more than one trait, regardless of the interacting partner. For example, SNP1 might interact with SNP2 to influence ANA, but interact with SNP3 to influence ACA. SNP1 affects both traits through interactions, but by interacting with different partners. In this case, we keep SNP1, but discard SNPs 2 and 3. We searched in this way for SNPs that influenced all four traits as part of interactions, and then all three traits as part of interactions, and so on until we had selected 1500 SNPs that were maximally epistatic and pleiotropic.

### Combined Analysis of Pleiotropy and Epistasis (CAPE)

We identified genetic interactions between SNPs using the Combined Analysis of Pleiotropy and Epistasis (CAPE) [14, 15]. This method combines information across multiple traits to infer directed genetic interactions. Combining information across traits not only increases power to detect genetic interactions, but also allows inference of the allele-to-allele direction of the interaction, which can be either positive (enhancing) or negative (suppressing). We have applied this analysis to multiple model organisms including yeast [15], fly cell lines [22], and mice [23-26]. This is the first application to human genetics.

As a preliminary analysis, we tested whether SNPs previously associated with SSc could be recapitulated in this small cohort. We performed linear regression to associate each SNP with each of the four traits.

We then decomposed the autoantibody and lung traits to four orthogonal eigentraits (ETs) using singular value decomposition (SVD) (Fig 2). This step concentrates trait variance into composite ETs potentially improving the ability to map weak genetic effects that are otherwise distributed across traits. We analyzed the first two ETs, which explained 71% of the variance across all four phenotypes (Fig 2A). These ETs contrast each of the autoantibodies with the other autoantibody and the two lung traits in turn (Fig 1A).

We performed linear regression on pairs from the 1500 SNPs for each ET. To avoid testing SNPs in linkage disequilibrium, we did not test any SNP pairs whose Pearson correlation coefficient exceeded 0.5. Out of 1,125,750 possible pairs between the 1500 SNPs plus the covariate sex, 1,125,593 SNP pairs (> 99.9%) passed this criterion. Another measure we pursued to false positive SNP pairs, was to test for population structure. Population structure, caused by heterogeneous relatedness among individuals in a cohort, can artificially inflate the association between SNPs and traits. To test whether population structure was influencing our association tests, we compared the quantile normalized distribution of the -*log*_10_(*p*) from association tests with each trait to the distribution of quantile normalized *p* values under the null hypothesis (Supplementary Fig 1). There was no systematic inflation of the estimates of significance. We therefore did not correct for population structure in this study.

We used the regression coefficients to calculate CAPE parameters that recast pair-wise epistasis as allele-to-allele influences that, in turn, modify allele-to-phenotype main effects. This process is described in detail elsewhere [14, 15], and we describe it briefly here. The first step in calculating CAPE coefficients is a reparametrization of the regression coefficients to obtain delta parameters (*δ*_1_ and *δ*_2_) which describe the degree to which the presence of one SNP influences the phenotypic effects of the other SNP:

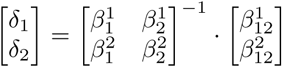

These parameters are then translated into directional (*m*_12_ and *m*_21_) coefficients that self-consistently describe how the two SNPs influence one another:

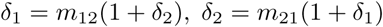

Standard errors for these interaction terms were estimated by propagating standard error terms from least squares regression with second-order Taylor expansion on the regression parameters [15].

Significance of the resulting model parameters were estimated through permutations with family-wise error rate estimation. We generated a null distribution as follows: ETs were randomly re-ordered relative to the genotypes, and we performed the single SNP scan, and the pairwise scan as with the original data. We repeated this process until we had generated a null distribution with 1.5 million values. We corrected empirical P values using false discovery rate (FDR) [27] and used a significance threshold of *q* < 0.05.

### Assignment of SNPs to genes

To assign each SNP in the final network to a gene, we downloaded SNP annotations from SNP Nexus [28-30] and assigned each gene to the nearest or containing gene.

### Coexpression analysis

We used the Search-Based Exploration of Expression Compendium (SEEK) [31] to identify genes that were co-regulated with genes identified by CAPE. SEEK is a web tool (http://seek.princeton.edu) that searches across thousands of gene expression data sets to find genes that are co-expressed with a user-defined query gene or gene set. The search can be restricted to data sets in individual tissues or that have associated keywords, thus providing context for the co-regulation.

Based on the CAPE results, we analyzed the two gene pairs: *WNT5A* and *RBMS3* in one query and *RBMS3* and *MSI2* in another. Using SEEK, we searched for co-regulated genes across all expression data sets. We also restricted our search to subsets of data sets to identify co-regulated genes in particular contexts (Table 2). For example, to look for tissue-specific co-regulation, we searched for co-regulated genes in all lung tissue data sets and all skin tissue data sets. Both tissues are relevant to SSc pathogenesis. However, in this study, we identified genetic interactions associated with lung function. We thus expected that genes co-regulated with the query genes in lung would better reflect the function of each interaction in this study. In addition to tissue-specific searches, we also searched for genes co-regulated with our query genes in disease-specific data sets. To perform these searches, we restricted the search to data sets associated with key words, for example autoimmunity and fibrosis.

In each of these queries, we identified the genes whose expression was significantly correlated with the pair of query genes (*p* < 0.05). These gene sets represent genes that are co-regulated with the query gene set across multiple tissues and disease contexts. We propose that functional enrichments of these gene sets are related to the function of the query gene pair in this disease cohort. To identify functional enrichment of each gene set, we used gProfileR [35] (See workflow).

Among the significantly enriched terms, were many that are known to be related to SSc, for example “extracellular matrix” and “wound healing.” We were particularly interested in enrichment for these terms that are known to be SSc-related and wanted to compare enrichment in SSc-related terms across the different SEEK queries. To do this, we looked only at enrichments of terms containing the following strings: “collagen”, “matrix”, “fibroblast”, “growth factor”, “vascular”, “vaso”, “wound”, and “wnt.” This is not an exhaustive list of the terms that are SSc-related, but is sufficient to to highlight major differences between functional enrichments of the two gene pairs across different contexts.

### Data Availability

Genotype and phenotype data are available from the Database of Genotypes and Phenotypes (DbGaP) (https://www.ncbi.nlm.nih.gov/gap) Accession: phs000357.v2.p1. Gene expression data are available from the Gene Expression Omnibus (GEO) (https://www.ncbi.nlm.nih.gov/geo/) Accession: GSE76808 A complete workflow including all code to generate results and figures is available here: https://github.com/annaLtyler/SSc_Epistasis_Workflow

## Supporting information

### S1 Fig. QQ plots of p values

To test for population structure in this patient cohort, we performed marker regression tests for each phenotype using sex as a covariate. We plotted the quantile normalized *p* values against the theoretical normal distribution. If population structure were affecting the SNP associations, we would expect to see systematic deviation from the line y = x in these plots. Rather we see that the distribution almost perfectly matches the theoretical distribution. The only deviations are a few significant p values in the HLA region for the ACA phenotype. We concluded that there was no population structure in this population and did not use a correction for population structure in our analysis.

### S2 Fig. Functional enrichment of co-expressed genes

We searched across multiple subsets of data sets in the Gene Expression Omnibus (GEO) for genes significantly co-expressed with either *RBMS3* and *WNT5A* or *RBMS3* and *MSI2*. This series of figures reports all terms significantly associated with each pair in each data subset. Bars show the -*log*_10_(*p*) of the enrichment. Blue highlighting indicates a term matching one of the key words we used to identify SSc-related terms.

## Acknowledgments

Genotype and phenotype data for the Genome-Wide Association Study in Systemic Sclerosis Study were provided by Maureen D. Mayes, University of Texas Health Science Center, Houston, Texas. Funding support for the original study was provided by the National Institutes of Health, and by other sources detailed in Nat Genet. 2010 May;42(5):426-9. Genome-wide association study of systemic sclerosis identifies CD247 as a new susceptibility locus. Radstake TR, Gorlova O, Rueda B, Martin JE, Alizadeh BZ, Palomino-Morales R, Coenen MJ, Vonk MC, Voskuyl AE, Schuerwegh AJ, Broen JC, van Riel PL, van ’t Slot R, Italiaander A, Ophoff RA, Riemekasten G, Hunzelmann N, Simeon CP, Ortego-Centeno N, Gonzàlez-Gay MA, Gonzàlez-Escribano MF; Spanish Scleroderma Group, Airo P, van Laar J, Herrick A, Worthington J, Hesselstrand R, Smith V, de Keyser F, Houssiau F, Chee MM, Madhok R, Shiels P, Westhovens R, Kreuter A, Kiener H, de Baere E, Witte T, Padykov L, Klareskog L, Beretta L, Scorza R, Lie BA, Hoffmann-Vold AM, Carreira P, Varga J, Hinchcliff M, Gregersen PK, Lee AT, Ying J, Han Y, Weng SF, Amos CI, Wigley FM, Hummers L, Nelson JL, Agarwal SK, Assassi S, Gourh P, Tan FK, Koeleman BP, Arnett FC, Martin J, Mayes MD.

